# Development of cell-free platforms for discovering, characterizing, and engineering post-translational modifications

**DOI:** 10.1101/2024.03.25.586624

**Authors:** Derek A. Wong, Zachary M. Shaver, Maria D. Cabezas, Martin Daniel-Ivad, Katherine F. Warfel, Deepali V. Prasanna, Sarah E. Sobol, Regina Fernandez, Robert Nicol, Matthew P. DeLisa, Emily P. Balskus, Ashty S. Karim, Michael C. Jewett

## Abstract

Post-translational modifications (PTMs) are important for the stability and function of many therapeutic proteins and peptides. Current methods for studying and engineering PTM installing proteins often suffer from low-throughput experimental techniques. Here we describe a generalizable, *in vitro* workflow coupling cell-free protein synthesis (CFPS) with AlphaLISA for the rapid expression and testing of PTM installing proteins. We apply our workflow to two representative classes of peptide and protein therapeutics: ribosomally synthesized and post-translationally modified peptides (RiPPs) and conjugate vaccines. First, we demonstrate how our workflow can be used to characterize the binding activity of RiPP recognition elements, an important first step in RiPP biosynthesis, and be integrated into a biodiscovery pipeline for computationally predicted RiPP products. Then, we adapt our workflow to study and engineer oligosaccharyltransferases (OSTs) involved in conjugate vaccine production, enabling the identification of mutant OSTs and sites within a carrier protein that enable high efficiency production of conjugate vaccines. In total, we expect that our workflow will accelerate design-build-test cycles for engineering PTMs.

## Introduction

Protein and peptide-based biologics play an important role in both treating and preventing a wide variety of illnesses. Currently, about 30% of all new US Food and Drug Administration (FDA) approved therapeutics entering the clinical setting are biologics.^1^ Common protein-based therapeutics include antibodies^2^, blood coagulants^3,4^ and protein-based vaccines^5,6^ among others^7^. Peptide drugs continue to mature as important options for treating microbial infection^8^ and diabetes^9^, among other conditions^10^, with over 800 new peptide therapeutics either in clinical development or undergoing preclinical studies^11^. Understanding how to design and produce protein and peptide-based therapeutics with optimal characteristics will continue to be a major focus in biological research.

For many biologics, post-translational modifications (PTMs) are important for the molecule’s overall stability and activity. Examples of PTMs include γ-carboxylation^12^, β-hydroxylation^12^, glycosylation^13^, and sulfation^14^, among many others^15^. Unfortunately, workflows for studying PTMs are often low throughput. For example, studies screening libraries of PTM installing enzymes or protein substrates often require overexpression of each variant in individual strains and labor-intensive protein purification steps. These methods are then coupled with low-throughput analytical methods such as mass spectrometry^16–18^, Western blotting^19^ or ELISA^20^, which are often time intensive or involve complex data analysis, severely limiting the throughput of sample analysis. Additionally, techniques used to directly measure interactions between PTM installing enzymes and their substrates, such as fluorescence polarization^21^, co-crystallization of the substrate in the enzyme active site^22,23^, and isothermal thermal calorimetry (ITC)^24^ are even lower-throughput, often limiting studies to single digits or tens of variants.

Advances in cell-free expression (CFE) systems have enabled the parallelized expression of proteins and peptides, which can facilitate the rapid characterization and engineering of PTMs. CFE systems^25–27^ involve using the transcription and translation machinery in crude extracts, rather than living cells, for protein synthesis. By supplementing the reaction with additional cofactors, an energy source, and salts, as well as a DNA template of the desired protein, many proteins can be produced in parallel in hours, alleviating the initial bottleneck of expressing large libraries of protein or peptide variants. Indeed, CFE systems have been successfully applied to a variety of high-throughput bioengineering applications, such as transcription factor engineering^28,29^, constructing metabolic^30^ and glycosylation pathways^31,32^, and studying substrate promiscuity of various PTM installing enzymes^33,34^. However, to date, many of these applications continue to rely on liquid chromatography and mass spectrometry-based approaches or the ability to connect the targeted protein function with a visual output such as sfGFP production. To take full advantage of the capabilities of CFE systems, there is a need for a generalized workflow for studying protein-peptide interactions and monitoring the installation of PTMs.

Here, we describe an *in vitro*, plate-based platform for characterizing and engineering PTMs by coupling cell-free protein synthesis (CFPS) with AlphaLISA. To showcase the utility of our workflow, we applied our method to two model systems: ribosomally synthesized and post-translationally modified peptides (RiPPs) and conjugate vaccines. To begin, we show that our workflow can be used to detect interactions between RiPP recognition elements and their native precursor peptides, a key first step in the biosynthesis of many RiPP products^35^. We then demonstrate how our workflow can be used to rapidly characterize which peptide residues are important for RRE binding as well as be integrated with computational prediction tools to create a biodiscovery pipeline for novel RiPP products. To show how our workflow can be used to directly measure the installation of PTMs, we then use the workflow to express and measure the activity of PglB from the organism *Campylobacter jejuni*^36^, an oligosaccharyltransferase (OST) with potential in conjugate vaccine production. Using a semi-rational approach, we design a library of 285 unique enzyme variants and use our cell-free workflow to identify 7 high-performing mutants, including a single mutant with a 1.71 fold improvement of glycosylation with a clinically relevant glycan. Finally, we demonstrate how our workflow can be used in the design of next generation conjugate vaccines by systematically scanning a carrier protein for sites accessible to *in vitro* glycosylation by PglB. In total, we expect that our workflow will accelerate the characterization and engineering of PTMs important for protein and peptide-based therapeutics.

## Results

Key to our workflow’s development was selecting a downstream analytical technique that could match the throughput expression capabilities of CFPS. AlphaLISA^37^ is an in-solution, bead-based assay version of ELISA that is amenable to acoustic liquid handling robots and small (1-2 μL) reaction sizes in 384- or 1536-well plate formats and has previously been used with cell-free systems to assess protein-protein interactions^38–40^. By requiring only liquid transfer and incubation steps, AlphaLISA facilitates the analysis of hundreds of reactions in hours.

### A cell-free, AlphaLISA based workflow can detect RRE-peptide interactions

As a model system for studying interactions between PTM installing proteins and their peptide substrate, we chose to work with ribosomally synthesized and post-translationally modified peptides (RiPPs) due to growing interest in their use as antimicrobial therapeutics^41–45^. Composed of an amino acid backbone, RiPPs biosynthetically originate as a precursor peptide composed of an N-terminal leader sequence and C-terminal core sequence^46^. Tailoring enzymes encoded within the same biosynthetic gene cluster (BGC) as the precursor peptide recognize a portion of the leader sequence and install post-translational modifications on the core sequence, producing the mature RiPP^46^. In over 50% of RiPP BGCs, the recognition of the leader sequence by tailoring enzymes is facilitated by a standalone protein or portion of a fusion protein containing a RiPP precursor peptide recognition element (RRE)^47^. In the absence of the RRE, individual reactions catalyzed by the tailoring enzymes often suffer from slow kinetics and low conversion rates^48^. Yet, despite their importance in catalyzing RiPP formation, current methods for studying interactions between RREs and their peptide substrate are low-throughput (e.g. fluorescence polarization^47,49,50^ and co-crystallization^22^), limiting studies to only single digits or tens of interactions at a time.

To begin, we selected a panel of 13 RREs from a range of RiPP classes and tested their expression in PURE*frex* via incorporation of FluoroTect^TM^ Green_Lys_ fluorescently labeled lysine (**Supplementary Fig.1**). For 9 of these proteins, we tested the native sequence as well as fusion proteins in which the predicted RRE domain of the protein was fused either N-terminally or C-terminally to maltose-binding protein (MBP) due to their size and/or origin from a radical *S*-adenosyl-L-methionine (SAM) enzyme potentially making PURE*frex* expression difficult. While some of the full-length constructs did produce soluble protein, we generally saw better expression when constructs were fused to MBP.

Encouraged by these results, we then tested the functionality of these RRE-containing proteins in an AlphaLISA based reaction with each of the RREs’ respective peptide substrates. We first expressed MBP-tagged RRE fusion proteins and N-terminally tagged sFLAG peptide substrates in individual PURE*frex* reactions (**Fig. 1a**). We then assayed for RRE-peptide recognition by mixing an RRE protein-expressing PURE*frex* reaction and the corresponding peptide substrate-expressing reaction with anti-FLAG donor beads and anti-MBP acceptor beads. Only in instances in which the RRE binds the peptide will the acceptor and donor bead be brought within close enough proximity to produce a chemiluminescent signal. A cross-titration of four different RRE-peptide pairs (PqqD, TbiB1, HcaF, TbtF) across multiple dilutions revealed a clear binding pattern consistent with RRE-peptide engagement (**Fig. 1b-1e**), which we do not observe when assaying MBP only with the respective peptides (**Supplementary Fig. 2**). Two of the pairs utilize fusion proteins containing only the predicted RRE domain (TbtF and HcaF), suggesting that predicted RRE domains, rather than full-length proteins, can be assessed for binding in this workflow.

**Figure 1.**
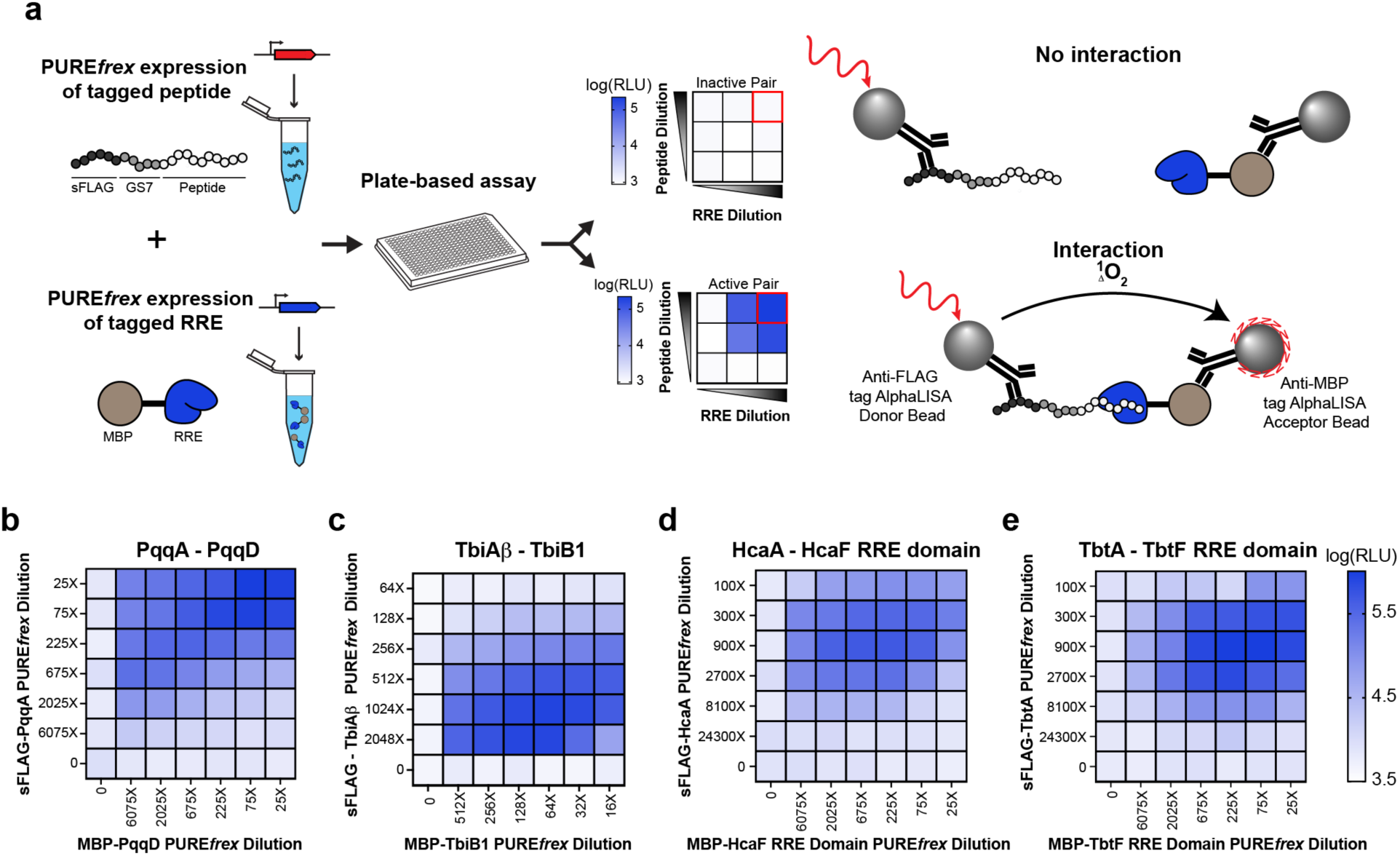
A cell-free, plate-based assay for detecting RRE-peptide interactions. (a) Schematic of the cell-free workflow. sFLAG-tagged peptides and MBP-tagged RREs are expressed in individual PURE*frex* reactions, mixed in a 384 well plate, and incubated to enable binding interactions. Addition of anti-FLAG AlphaLISA donor beads and anti-MBP AlphaLISA acceptor beads enables detection of interactions between the RRE and peptide of interest. PURE*frex* reactions of precursor peptide and RRE for (b) pyrroloquinoline quinone (PQQ), (c) a putative lasso peptide from *Thermobacculum terrenum* ATCC BAA-798, (d) a heterocycloanthracin from *Bacillus* sp. Al Hakam, and (e) thiomuracin, a thiopeptide from *Thermobispora bispora* were cross-titrated across different dilutions and assessed for binding interactions via AlphaLISA. Data are representative of at least three biological replicates.

### Cell-free workflow enables mapping of residues important for RRE-peptide binding

We next sought to benchmark our workflow’s ability to identify peptide residues important for RRE binding. The current state of the art for this type of study involves competition fluorescence polarization assays with an alanine scan library of peptide variants^51^. To test our methodology, we characterized binding of the RRE domain of TbtF, the cyclodehydratase involved in thiomuracin^51,52^ biosynthesis, to the leader sequence of TbtA (**Fig. 2a**). Previous work by Zhang et al. found that mutating residues L(-32), L(-29), M(-27), D(-26), and F(-24) to an alanine resulted in the greatest reduction of binding affinity by TbtF^51^, a result that our workflow confirms as evidenced by a greater than 100-fold decrease in AlphaLISA signal compared to the wild-type peptide sequence. Interestingly, our workflow also found that the mutation D(-30)A also resulted in a greater than 100-fold decrease in AlphaLISA signal whereas Zhang et al. observed only a small reduction in binding affinity using the same peptide variant in their assay. Regardless, by utilizing simple DNA manipulation techniques, we were able to construct the library of DNA templates, express peptide and protein constructs, and assay for binding activity within hours. This is faster than conventional cloning, transformation, expression, and purification workflows required for fluorescence polarization competition assays, which can take weeks to months.

**Figure 2.**
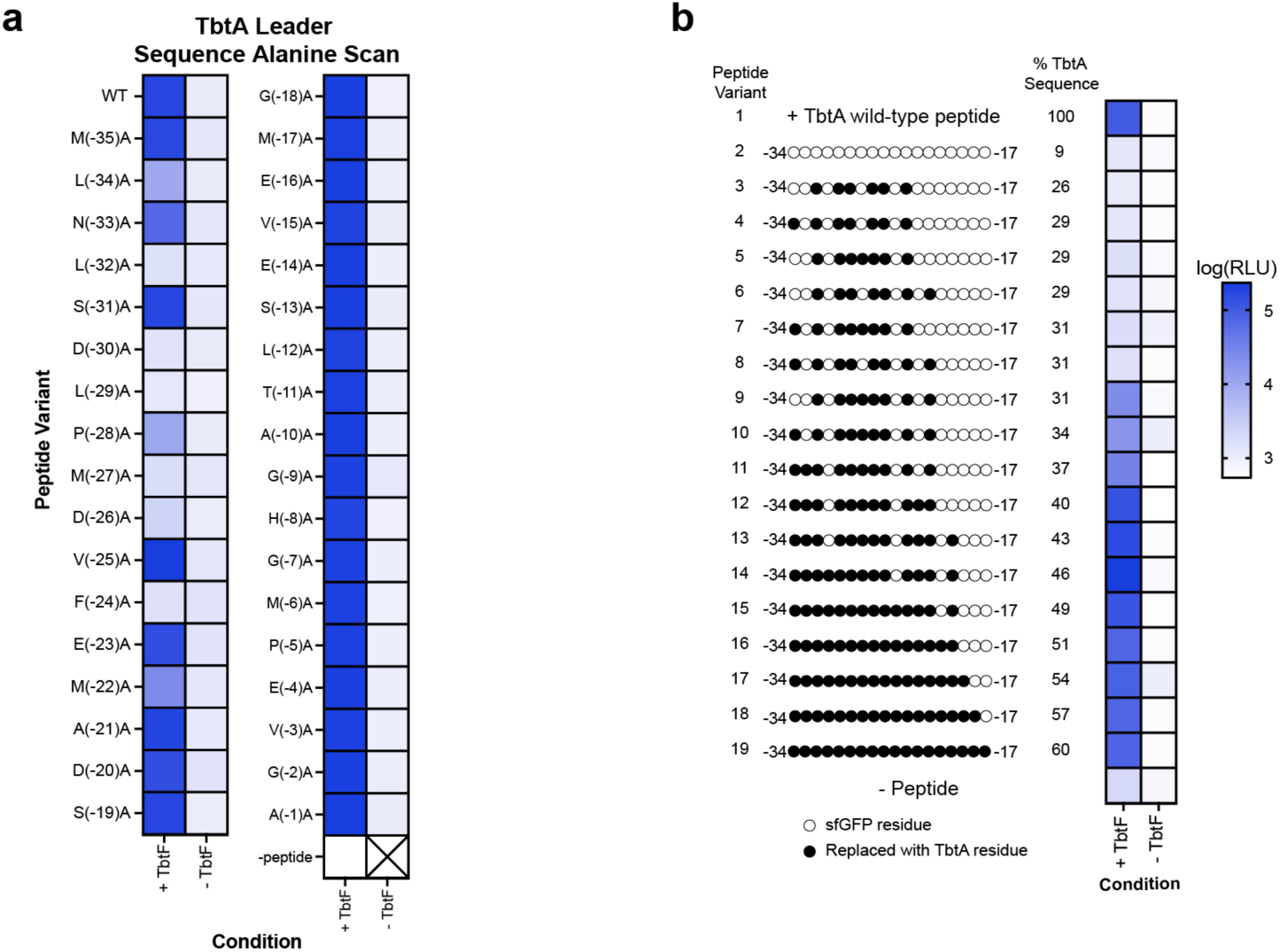
Cell-free workflow identifies peptide residues important for binding by TbtF. (a) An alanine scan library of the leader sequence of TbtA was expressed in individual PURE*frex* reactions and assessed for binding interactions in the presence of MBP-TbtF RRE domain using AlphaLISA. (b) A synthetic peptide library was constructed using the first 40 amino acids of sfGFP. Variants of the sfGFP were then constructed by replacing residues in the peptide identified in the alanine scan as important for binding by TbtF with the corresponding residue in the wild-type TbtA leader sequence. Each peptide variant was expressed in an individual PURE*frex* reaction, and then assessed for binding interactions in the presence and absence of TbtF using AlphaLISA. Peptide variant 2 contains 9% identity to TbtA wild-type peptide due to sharing residues, G(-2), G(-7) and G(-9). For simplicity, only amino acids between the -34 and -17 position are depicted, however each peptide was composed of 40 amino acids reflecting the length of the TbtA leader sequence with an additional 5 amino acid linker. Sequences for each of the peptide variants assayed in panel b are provided in **Supplementary Table 1**. All data are presented as the mean of n = 3 technical replicates.

We next asked whether we could use the characterized peptide-binding landscape to inform the design of a synthetic peptide capable of binding to TbtF. As a starting point, we constructed a synthetic peptide sequence the same length as the leader sequence of TbtA that does not bind to TbtF (**Fig. 2b**; peptide variant 2), using the first 40 amino acids of sfGFP with a G(-18)T mutation to ensure all residues in the region of interest differed from the wild-type TbtA leader sequence. We then created peptide variants by replacing residues in the synthetic peptide with residues identified from the alanine scan as important for binding by TbtF, starting with the six residues (L(-32), D(-30), L(-29), M(-27), D(-26), and F(-24) that when mutated to an alanine resulted in the greatest decrease in AlphaLISA signal. We were unable to detect binding interactions between this engineered peptide variant (peptide variant 3) and TbtF. Next, we created peptide variants 4-10 by adding individually, or in combination, residues L(-34), P(-28), and M(-22), which in our initial screen also appeared to slightly reduce binding affinity to TbtF when mutated to an alanine. Adding both P(-28) and M(-22) (peptide variant 9) to peptide variant 3 enabled weak binding by TbtF, with ∼25% AlphaLISA signal of the wild-type TbtA leader sequence. Further addition of residues resulted in a synthetic peptide (peptide variant 12) that is 40% identical to the leader sequence of TbtA (L(-34), N(-33), L(-32), D(-30), L(-29), P(-28), M(-27), D(-26), F(-24), E(-23), and M(-22) and exhibits binding to TbtF (AlphaLISA signal) that is approximately equal to that observed with the wild-type TbtA leader sequence peptide. Interestingly, adding residues D(-20) and S(-31) (peptide variant 14) increased the signal further to ∼2-fold higher than that observed with the wild-type TbtA leader sequence. These results highlight our assay’s ability to rapidly identify specific residues involved in RRE-peptide binding interactions and design new peptide sequences with the minimum number of residues required for RRE engagement.

### Cell-free workflow as a screening tool for *in vitro* activity of computationally identified RRE-peptide pairs

Successful heterologous expression of computationally predicted RiPP products *in vivo* remains a challenge due to the inability to precisely control the timing and extent to which each BGC protein is expressed as well as the presence of necessary cofactors^53^. As a result, there is increasing interest in using *in vitro* based approaches to prototype computationally predicted RiPP BGCs^53^. Due to the importance of RREs in RiPP biosynthesis, we asked whether our workflow could be used to characterize RRE binding for uncharacterized lasso peptide BGCs computationally predicted via AntiSMASH^54^. Lasso peptides are a class of high interest RiPPs due to their unique lariat structure which imparts the molecule with a wide range of beneficial characteristics, such as heat and protease stability^55,56^. Additionally, lasso peptides have displayed a variety of bioactivities, including antimicrobial activity^57–62^. Biosynthetically, lasso peptide BGCs typically encode (i) a precursor peptide, (ii) an RRE and (iii) a protease, or a fusion protein encoding both the RRE and protease, as well as (iv) a cyclase^55^. In all reported lasso peptide BGCs, RREs are important for guiding the protease to the precursor peptide substrate and in some cases is also required for cyclization by the cyclase^24,63–67^. Previous works have also successfully utilized cell-free systems to study lasso peptide biosynthesis^68^.

To begin, we used AntiSMASH^54^ to identify a total of 2,574 lasso peptide BGCs from a collection of 39,311 diverse genomes (**Fig. 3a, Supplementary Table 2**). Of these, 1,882 BGCs were predicted to contain a complete collection of essential lasso peptide biosynthetic enzymes (**Supplementary Table 3**). With an eye towards discovering lasso peptides, we compared the identified BGCs to known lasso peptides by constructing a sequence similarity network of the predicted core peptide sequences and annotating known sequences within the resulting network. Sequences that matched computationally predicted but not experimentally verified sequences reported in the literature were maintained in the dataset while those that had been produced experimentally were removed; in doing so, we reasoned that our workflow could help assess the accuracy of predictions generated by others in the field while also ensuring novelty of any clusters successfully reconstituted. From the remaining predicted BGCs, 47 were selected for study from 32 unique genera based on their potential for therapeutic relevance, namely antibiotic activity (**Extended Data Set**).

**Figure 3.**
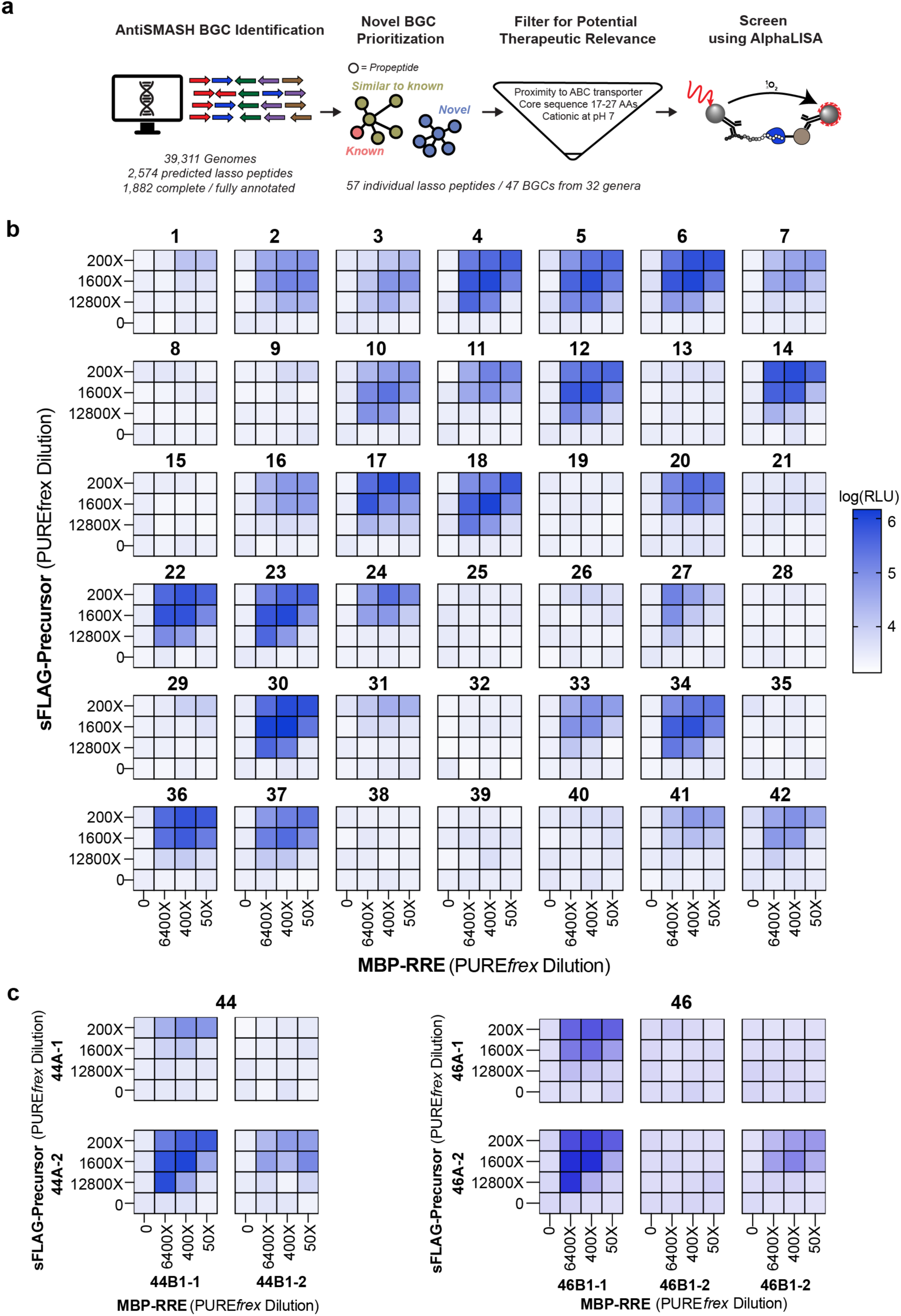
Computational guided screen of lasso peptide RREs. (a) Overall prediction and screening workflow for lasso peptide BGCs. (b) For predicted BGCs with a single RRE and precursor peptide, individual PURE*frex* reactions expressing RREs and respective peptide were cross-titrated and assessed for binding activity via AlphaLISA. (c) For predicted biosynthetic gene clusters with multiple RREs and precursor peptides, all possible combinations of RRE and precursor peptide were assessed for binding activity via AlphaLISA (select combinations shown, see **Supplementary Figure 5** for additional pairwise combinations. All data shown are single replicate, with confirmation reactions performed in biological triplicate provided in **Supplementary Figure 6**.

Of the 47 predicted lasso peptide BGCs, 5 were predicted to contain more than one precursor peptide and/or RRE, bringing the total number of predictions to 57 unique precursor peptides and 52 unique RREs. We applied our cell-free workflow to screen all 57 predicted precursor peptides with their associated predicted RREs (**Fig. 3b** and **3c**; **Supplementary Fig. 6**). To account for potential differences in expression levels in the PURE*frex* reactions as well as the fact that RREs reported in literature have a range of binding affinities, we tested each peptide-RRE pair at multiple concentrations. In instances where multiple RREs or precursor peptides were predicted in the same BGC, we screened all pairwise combinations. In total, we screened 72 different RRE-peptide pairs, 42 RRE-peptide pairs from clusters with a single predicted RRE and peptide pair (**Fig. 3b**) and 30 different combinations of RRE and peptides from clusters with multiple predicted genes for each (**Fig. 3c** and **Supplementary Fig. 6**).

Our initial screen yielded clear binding patterns for 27 of the 42 individual RRE-peptide pairs and 24 of the 30 RRE-peptide combinations from larger clusters (**Fig. 3b**, **Fig. 3c**, and **Supplementary Fig. 6**). A subsequent validation experiment assaying all RRE-peptide pairs in biological triplicate at the dilution condition that yielded the highest AlphaLISA signal confirmed the results of our screen, with RRE-peptide pairs that produced higher AlphaLISA signal in our initial screen generally producing higher AlphaLISA signal in the validation experiment (**Supplementary Fig. 7**). Notably, in addition to identifying functional RREs and peptide pairs, our methodology enables the rapid interrogation of more complex clusters. For example, 44B1-1 can bind to both precursor peptides identified in the cluster while 444B1-2 can only bind to the second precursor peptide (**Fig. 3c**). Similar behavior emerged in cluster 46 in which predicted RREs bound to both, only one, or neither of the precursor peptides (**Fig. 3c**).

Using the results from our large-scale RRE screen, we prioritized clusters identified as “hits” for complete biosynthesis of a mature lasso peptide *in vitro*. To do so, we expressed precursor peptides in PURE*frex* reactions and purified each related tailoring enzyme heterologously expressed in *E. coli.* Small scale (10 μL) reactions were assembled by combining precursor peptides and purified tailoring enzymes and analyzed via matrix-assisted laser desorption/ionization time-of-flight (MALDI-TOF) MS after overnight incubation at 37 °C. By testing several clusters, we successfully produced a peptide with the topology of a lasso peptide, from BGC 24 (**Supplementary Fig. 8**). Subsequent characterization experiments confirmed that the production of Las24 is time dependent (**Supplementary Fig. 9**), that each of the proteins in the predicted BGC is necessary for maturation (**Supplementary Fig. 10**), the structure of the molecule (**Supplementary Fig. 11**), and resistance to carboxypeptidase (a common confirmation of threaded topology) (**Supplementary Fig. 12**). As this work was ongoing, King et al. reported the heterologous production of a lasso peptide (termed Las-1010) in *E. coli* from the same biosynthetic cluster we report as Las24^69^. In addition to demonstrating *in vivo* biosynthesis of Las-1010, it was found that Las-1010 exhibits weak antibacterial activity against some bacterial strains^69^. While we are not the first to produce Las-1010, our ability to independently produce Las-1010 in an *in vitro* setting confirms the utility of our workflow for prototyping RiPP BGCs in cell-free systems.

### A cell-free, AlphaLISA based workflow for prototyping *in vitro* glycosylation reactions

As a model system for synthetic engineering of post-translational modifications, we asked whether a modified version of our cell-free AlphaLISA workflow could be used to monitor glycosylation activity of OSTs with bacterial glycans relevant to conjugate vaccine production. Conjugate vaccines, composed of a pathogen specific polysaccharide antigen (such as an O-antigen polysaccharide or capsular polysaccharide (CPS)) linked to an immunogenic carrier protein, are a promising strategy to protect against bacterial infections^70^. Both the glycan and carrier protein play an important role in developing long-lasting immunity; the glycan acts as the antigen and trains the immune system to recognize glycans that are natively displayed on the outside of pathogenic bacteria while the carrier protein enables the induction of a T-cell dependent immune response, leading to a stronger and longer lasting immune response compared to immunization with glycan alone^71^.

One of the current challenges with manufacturing conjugate vaccines is the reliance on a multi-step process in which the glycan is isolated from the targeted pathogenic bacteria and chemically conjugated to a separately produced carrier protein^72^. These processes require specialized equipment, stringent purification steps to remove the pathogenic bacteria, and can result in a heterogeneous final product. To address these challenges, recent work in the field of glycobiology and synthetic biology have developed alternative cell^73–78^ and cell-free^79–81^ based methods for producing conjugate vaccines using OSTs to site-specifically transfer glycans onto a carrier protein.

Previous work in the field has demonstrated that the CPS from *S. pneumoniae* serotype 4 can be transferred onto a protein *in vivo* by the OST PglB from the organism *Campylobacter jejuni* (*cj*PglB)^82^. To recapitulate these findings in an *in vitro* setting, we first expressed *cj*PglB in a CFPS reaction supplemented with nanodiscs, which act as a membrane mimic into which membrane-bound *cj*PglB can solubly express^83^. We subsequently mixed CFPS-expressed *cj*PglB with a separately expressed acceptor protein and a crude membrane fraction enriched with the CPS from *S. pneumoniae* serotype 4 (**Supplementary Fig. 13a**) in a process we term an *in vitro* glycosylation (IVG) reaction. By including a 6xHis tag on the acceptor protein, we can perform a Western blot to confirm transfer of the targeted bacterial glycan onto the protein, with the banding pattern above the aglycosylated protein corresponding to transfer of different chain lengths of the bacterial glycan (**Supplementary Fig. 13b**).

Having confirmed that we can achieve glycosylation in IVG reactions with the CPS from *S. pneumoniae* serotype 4, we then asked whether we could adopt our cell-free AlphaLISA based workflow to detect glycosylation (**Fig. 4a**). By incorporating an anti-glycan antiserum into the AlphaLISA reaction and using Protein A AlphaLISA donor beads and anti-6xHis AlphaLISA acceptor beads, we hypothesized that we would be able to distinguish between glycosylated and aglycosylated proteins. Indeed, when we prepared IVG reactions using an acceptor protein containing a sequon (a short stretch of amino acids) that can (DQNAT) or cannot (AQNAT) be glycosylated by *cj*PglB and analyze the reactions using AlphaLISA, we observe a distinct banding pattern only when we use the protein containing DQNAT, confirming our ability to discriminate between glycosylated and aglycosylated samples (**Fig. 4b & Fig. 4c**).

**Figure 4.**
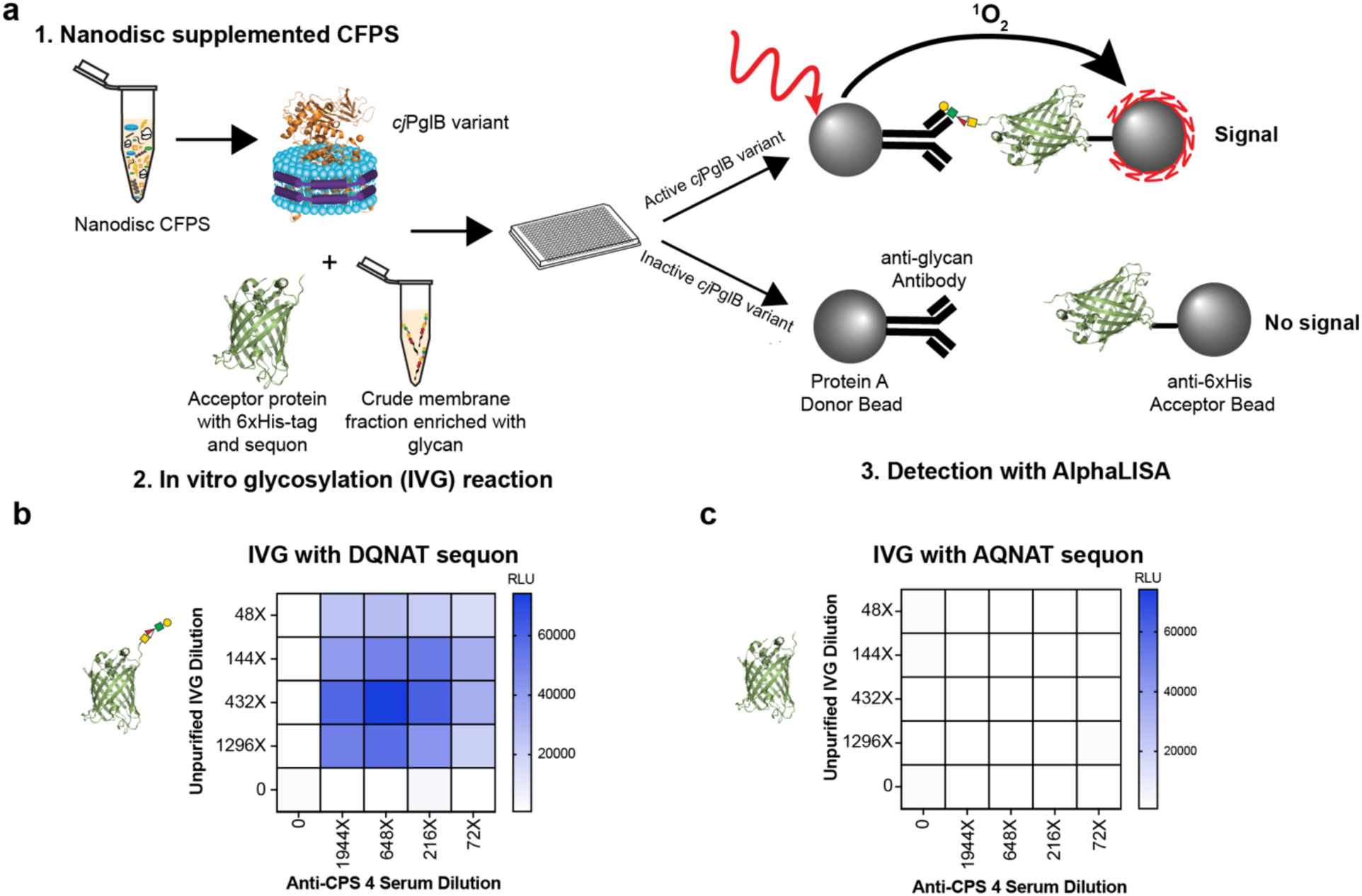
CFPS and AlphaLISA can be combined to prototype *in vitro* glycosylation reactions. (a) Schematic of the cell-free workflow. Nanodisc supplemented CFPS reactions were first used to express *cj*PglB variants and then mixed with an acceptor protein containing a 6xHis tag and sequon and crude membrane fraction enriched with a bacterial glycan of interest. Samples were then analyzed using Protein A AlphaLISA donor beads and anti-6xHis AlphaLISA acceptor beads. (b) An IVG reaction using *cj*PglB, crude membrane fraction enriched with CPS from *S. pnuemoniae* serotype 4, and sfGFP with a 6xHis tag and either (b) a DQNAT sequon or (c) an AQNAT sequon was serially diluted, mixed with varying concentrations of *S. pneumoniae* CPS 4 antiserum, and analyzed via AlphaLISA. All data are representative of at least three independent experiments.

### Cell-free workflow enables semi-rational engineering of *cj*PglB for increased transfer efficiency of CPS from *S. pneumoniae* serotype 4

While *cj*PglB has demonstrated some glycan substrate promiscuity^73,75,77,79–81^, the efficiency with which it can glycosylate acceptor proteins with different glycans varies widely. To address this challenge, recent work has demonstrated that mutating PglB can lead to improvements in glycosylation efficiency^76,84^. When we tested two previously identified *cj*PglB mutants^76^,we observed improvements in glycosylation efficiency with CPS from *S. pneumoniae* serotype 4 (**Supplementary Fig. 13b**). However, due to the unique structure of each bacterial polysaccharide, we hypothesized that the optimal *cj*PglB mutation(s) required for high efficiency transfer likely differs from one glycan to another.

Towards this goal, we asked if our workflow could be used to express libraries of mutated *cj*PglB variants to screen for improved glycosylation efficiency with the CPS from *S. pneumoniae* serotype 4. To begin, we identified 15 *cj*PglB residues for site saturation mutagenesis, 9 residues (Y77, S80, S196, N311, Y462, H479, K522, G476, and G477) based on their predicted location within 4 angstroms of where the innermost sugar of the *cj*PglB native glycan sits within the *cj*PglB active site^76^, 3 residues (Q287, L288, and K289) within external loop 5 (EL5) which have previously been shown to be highly mutable^76^, and 3 additional residues (D475, K478, and L480) located in a flexible loop located directly above the nitrogen atom of the amide group on the acceptor protein where the glycan will be covalently linked^84^ (**Fig. 5a & 5b**). We then designed a library of *cj*PglB mutants in which each of these 15 residues was individually mutated to all 19 other amino acids, resulting in a library of 285 unique single mutant *cj*PglB constructs with 15 additional wild-type sequences.

**Figure 5.**
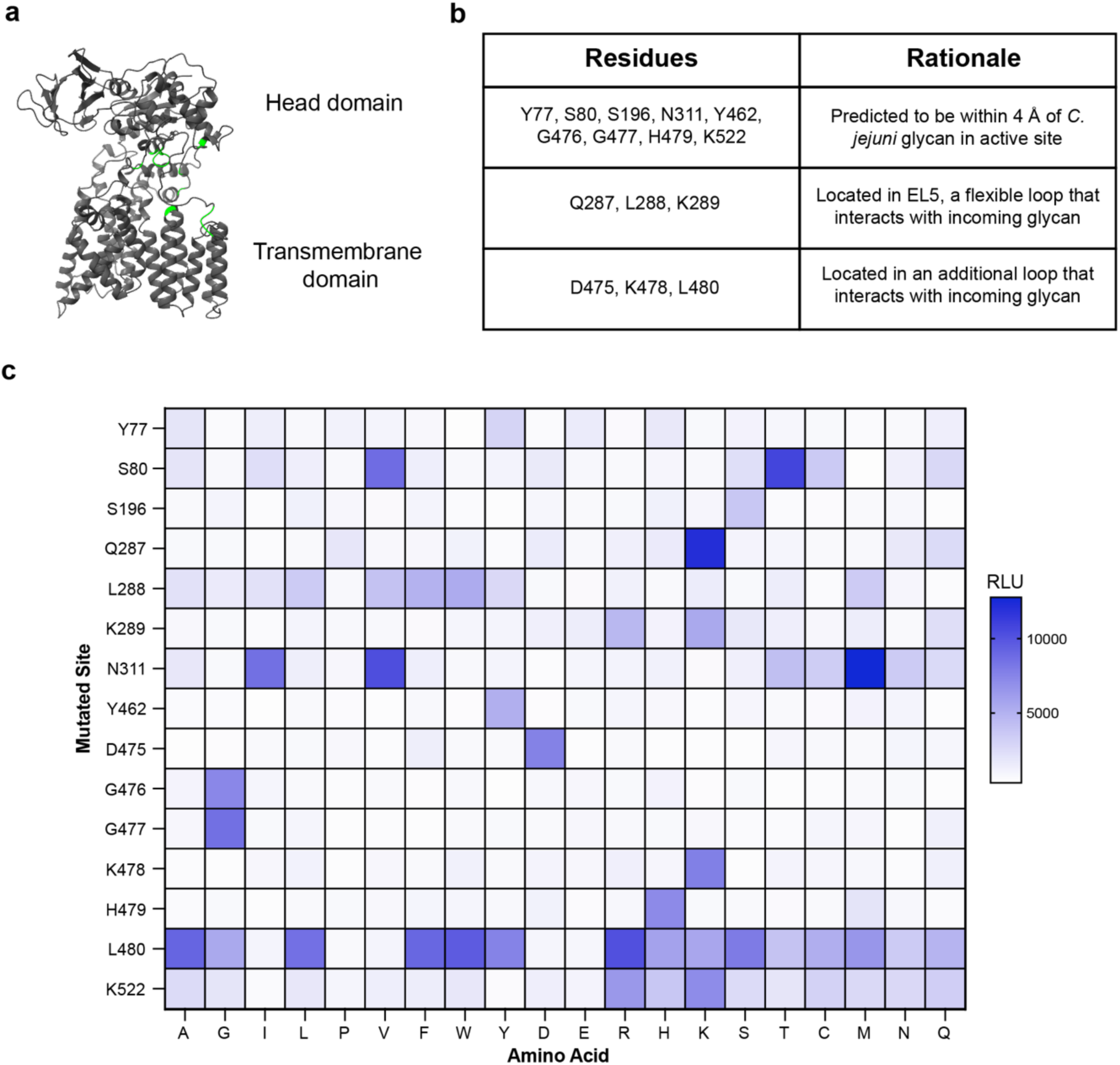
Cell-free workflow identifies high-efficiency *cj*PglB mutants. (a) Homology model of *cj*PglB^76^. Sites chosen for site saturation mutagenesis are highlighted in green. (b) Rationale of sites chosen for mutagenesis. (c) AlphaLISA results for IVG reactions containing crude membrane fraction enriched with CPS from *S. pnuemoniae* serotype 4, sfGFP with a 6xHis tag, and a unique *cj*PglB mutant. Data are a mean of AlphaLISA signals produced by duplicate IVG reactions.

Using our cell-free workflow, we then expressed the complete library of 300 constructs and assayed for activity (**Fig. 5c**). Ten *cj*PglB mutants (S80V, S80T, Q287K, N311I, N311V, N311M, L480A, L480W, and L480R) produced higher AlphaLISA signal than the WT *cj*PglB construct. Most sites were inflexible to mutation and produced no hits, whereas the sites S80, N311, and L480 produced multiple high-signal mutants. Control reactions that contained all reaction components except the *S. pneumoniae* serotype 4 anti-serum produced AlphaLISA signal equivalent to background (**Supplementary Figure 15a**), and duplicate measurements for each *cj*PglB mutant were consistent **(Supplementary Figure 15b**). To validate the results of our screen, we performed a Western blot of IVG reactions glycosylating the clinically-relevant carrier protein *Haemophilus influenzae* protein D (PD) with CPS from *S. pneumoniae* serotype 4 using each of the seven highest-signal mutants (S80V, S80T, Q287K, N311I, N311V, N311M, and L480R) and compared the transfer efficiency to WT *cj*PglB **(Supplementary Figure 16**). Each identified mutant produced a higher transfer efficiency than the WT enzyme, and PglB^Q287K^ raised the transfer efficiency by 38% (∼1.7x) to an efficiency of 91%.

### Cell-free workflow enables rapid identification of sites accessible for glycosylation in *in vitro* glycosylation reactions

The conventional technology to produce conjugate vaccines uses chemical methods to randomly conjugate glycans to a carrier protein^72^, which can modify T cell epitopes on the carrier protein and reduce vaccine immunogenicity^85,86^. In comparison, enzymatic production of conjugate vaccines using OSTs enables site-specific glycosylation of a carrier protein only at the synthetically inserted sequon, which could be used to avoid immune epitopes and lead to enhanced vaccine immunogenicity and reduction in the dose required to protect against each bacterial serotype in a multivalent vaccine^87,88^. To date, our cell-free approach to produce conjugate vaccines has relied on placing the sequon at the C-terminus of carrier proteins^79–81^. Since glycan placement may impact vaccine immunogenicity, we used our high-throughput cell-free workflow to screen a library of unique carrier protein constructs in which a sequon was placed between every pair of amino acids throughout the carrier protein PD to discover novel sites within a clinically relevant carrier protein that can be enzymatically glycosylated *in vitro* to form *S. pneumoniae* serotype 4 conjugate vaccines (**Fig. 6a**).

**Figure 6.**
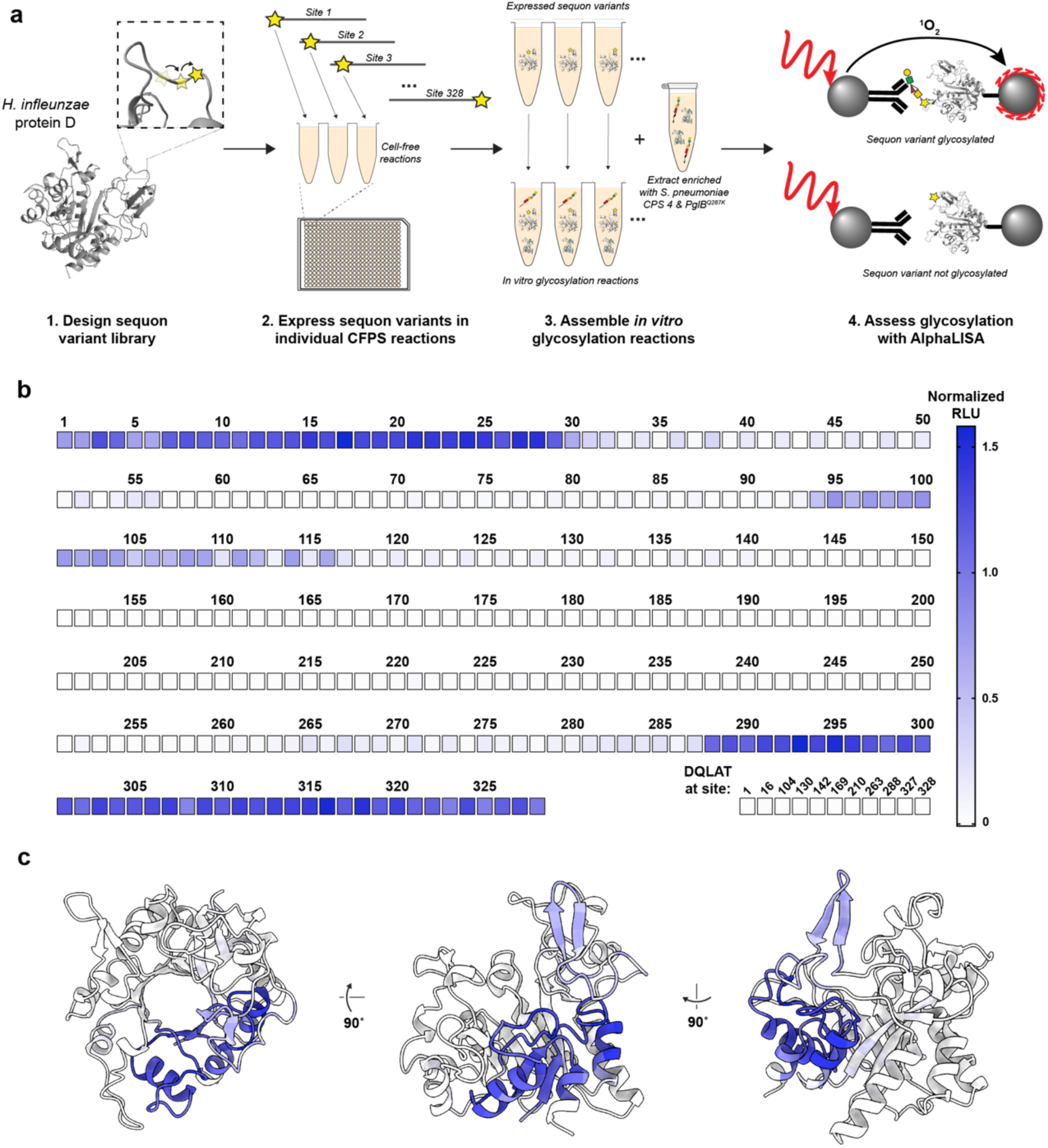
Sequon walkthrough of *H. influenzae* protein D. (a) Schematic of the cell-free workflow. A library of PD constructs was designed with a glycosylation sequon inserted between every two amino acids. Each sequon variant was expressed in an individiaul CFPS reaction. Following expression, sequon variants were combined with extract enriched with *S. pneumoniae* CPS 4 glycan and *cj*PglB^Q287K^ to form IVG reactions. IVG products were assessed for glycosylation with AlphaLISA. (b) AlphaLISA results for sequon walkthrough from N-terminus (site 1) through C-terminus (site 328). (c) AlphaLISA results from (b) mapped onto the crystal structure of PD.

In total, our library contained 328 unique PD sequences in which the glycosylation sequon “DQNAT,” surrounded by short linkers, was placed between every two amino acids in the carrier protein, beginning with an N-terminal sequon placement and ending with a C-terminal sequon placement. We then applied our cell-free workflow to this library by synthesizing each construct in a CFPS reaction, combining each synthesized carrier with CPS from *S. pneumoniae* serotype 4 and *cj*PglB^Q287K^ – the high efficiency mutant identified in the OST mutagenesis screen – and assessing glycosylation of each carrier protein construct in parallel in 1 µL AlphaLISA reactions using conditions optimized to detect low glycosylation levels **(Fig. 6b; Supplementary Figure 18**).

Our screen identified three sections of PD that were amenable to glycosylation in IVG reactions: 32 sites at the N-terminal end of the carrier sequence, a stretch of ∼20 internal sites, and 40 sites at the C-terminal end of the carrier sequence (**Fig. 6c & 6d**). Mapping AlphaLISA signal to a crystal structure of PD reveals one 3-dimensional section of the carrier protein that is able to be glycosylated (**Figure 6d**)^89^. In total, 94 sequon positions showed statistically significant signal above a negative control containing the sequon “DQLAT” (**Supplementary Figure 18a**), with top performing variants producing signal >100x above background. Triplicate measurements for each sequon variant were consistent with each other, and all negative control variants had signal equivalent to background (**Supplementary Figure 19a & 19b**). A selection of sequon variants with high and low AlphaLISA signal were validated via Western blot, confirming the accuracy of our workflow for comparing glycosylation efficiency of different carrier protein constructs (**Supplementary Figure 20**).

## Discussion

Here we developed an integrated workflow for expressing and characterizing proteins involved in PTM installation by combining methods for cell-free DNA assembly and amplification, cell-free protein synthesis, and binding characterization via AlphaLISA. We show that the platform is generalizable, fast (steps are carried out in hours), and readily scalable to 384- or 1536-well plates without the need for time intensive protein purification or cell-based cloning techniques. Moreover, the platform is designed with automation in mind, with each step consisting of simple liquid handling and temperature incubation steps. We showed the utility of the platform for characterizing the activity of both known and previously unidentified RREs involved in RiPP biosynthesis as well as towards engineering systems for efficient conjugate vaccine production.

Through our work characterizing the binding activity of TbtF to TbtA, we find that our methodology can within hours of obtaining DNA samples recapitulate findings obtained using traditional approaches that take days to weeks to perform. As a result, we can use our workflow for more advanced PTM engineering strategies. For example, recent work has created novel RiPP products by engineering peptide substrates to contain leader sequences recognized by tailoring enzymes from multiple classes of RiPPs^90^. In doing so, a peptide substrate was modified with RRE-dependent tailoring enzymes from two different BGCs. Creating more complex systems with even greater numbers of RRE-dependent modifications will require an understanding of appropriate design rules for enabling recognition of the precursor peptide by the desired tailoring enzymes. Using the information gained by mutational scanning, we were able to systematically produce a synthetic peptide with only 40% identity to the wild-type peptide that exhibits AlphaLISA binding signal on par with the wild-type peptide. Understanding the minimal set of amino acid residues required for recognition will be important for engineering increasingly complex molecules. Our workflow provides a method for understanding and prototyping these requirements.

Additionally, by coupling our workflow with computational prediction tools, we demonstrated how our platform can screen for natural product BGCs likely to function in an *in vitro* setting. While we were able to produce Las-1010 (Las24), many of the clusters we detected binding activity for in our screen did not ultimately lead to successful production of a mature lasso peptide. We hypothesize that this could be due to a number of reasons, including that the *in vitro* reaction environment may lack other important components natively included in *in vivo* systems, such as auxiliary genes as has been demonstrated for other natural products^91,92^. Regardless, by eliminating clusters with a non-functional first step (no RRE binding), we narrowed down the number of proteins needed to be expressed and purified for attempts at *in vitro* reconstitution. Using conjugate vaccines as a test system, we also demonstrated how our workflow can be used to rapidly engineer both enzymes and substrates for more efficient biomanufacturing systems. With our platform, we rapidly tested 285 unique mutants of *cj*PglB to enable efficient production of conjugate vaccines with the CPS from *S. pneumoniae* serotype 4, a major cause of pneumonia in disadvantaged communities^93^. Importantly, the top performing mutations discovered in our screen are different than those discovered in previous *cj*PglB mutagenesis experiments for other unrelated glycans, demonstrating the importance of a fast, high-throughput method to discover mutations unique for transferring each pathogen glycan of interest^76^. Because the identity of all tested mutants, including both low and high-performing mutations, is known at the time of assay, we believe our workflow could be readily interfaced with machine-learning guided strategies^94,95^ to more rapidly engineer oligosaccharyltransferases.

Towards the design of next generation conjugate vaccines, we used our workflow to rapidly assess the glycosylation of 328 unique variants of PD *in vitro* to discover novel, high efficiency glycosylation sites throughout the carrier protein. Over a quarter of all sites produced AlphaLISA signal significantly higher than a negative control. Mapping sites that produced high AlphaLISA signal to a crystal structure of PD reveals one section of the protein that was highly glycosylated, suggesting that steric effects may play a role in determining the ability to glycosylate unique sequon positions^96^ or that placing sequons in other locations of the protein may lead to low protein expression in CFPS reactions. Future work will focus on combining highly efficient engineered OSTs with identified glycosylation sites to create homogeneous vaccines with multiple glycan attachments per carrier, which may offer additional improvements to immunogenicity^85,97^.

One important limitation to our work is that the results we obtain are semi-quantitative. With our current platform design, we can compare the relative binding affinity of different RRE-peptide pairs or glycosylation efficiency of specific OST mutants but are unable to provide exact quantitative measurements of these phenomena (e.g. k_d_, % glycosylation, etc.). Thus, we suggest that our method can be integrated complementary to more traditional assays by first using our workflow as a screening tool to down-select specific protein variants for follow-up experiments.

In total, we developed a versatile platform for characterizing and engineering PTMs. We expect that this platform can be applied to other classes of PTMs and will accelerate the design and production of biologics with complex PTMs and improved therapeutic properties.

## Materials and Methods

### Methods for RiPPs work

#### DNA design and preparation

For the initial screen of known RRE’s, gene constructs were ordered from Twist Biosciences (synthesized into pJL1 backbone between NdeI and SalI restriction sites). Briefly, sequences were retrieved from literature or Uniprot and codon optimized using the IDT Codon Optimization Tool. For full length RRE constructs, a codon optimized sequence for a Twin-Strep tag and PAS11 linker were added to the N-terminus of the nucleotide sequence. MBP-fusion RRE constructs were constructed by replacing either the C-terminus (for proteins in which the RRE domain was predicted to occur in the N-terminus) or N-terminus (for proteins in which the RRE domain was predicted to occur in the C-terminus) portion of the sequence with codon optimized sequences for MBP and a GS7 linker. For precursor peptide sequences, sequences encoding either the full-length precursor or leader sequence were fused to an N-terminal sFLAG tag and GS7 linker.

For all peptide sequences used in AlphaLISA based assays, an N-terminal sFLAG tag and GS7 linker were incorporated into the design. For sequences utilized in the AlphaLISA alanine scan workflow, each alanine variant peptide was constructed by replacing the corresponding wild-type codon with “GCC”. To construct synthetic sfGFP peptides, the first 40 amino acids of sfGFP (with a G23T mutation) was first codon optimized. Each variant was then constructed by replacing the appropriate wild-type codon with the codon corresponding to the desired residue change. All peptide sequences were ordered as eBlocks with overhang to a linearized pJL1 backbone for use in Gibson Assembly reactions.

For all computationally predicted lasso peptide proteases and cyclases, the predicted gene sequences were codon optimized using the IDT Codon Optimization Tool. At the N-terminus of each sequence, maltose binding protein (MBP) and a short linker were incorporated to enable soluble expression and detection via AlphaLISA based assays. All genes were synthesized by Twist Biosciences either in pJL1 (for expression in PURE*frex*) or in a modified pET vector (for *in vivo* expression). The corresponding (untagged) precursor sequences were also synthesized by Twist Biosciences in pJL1 for use in assembling complete lasso peptide BGCs.

DNA templates for expression in PURE*frex* were prepared either in plasmid form using ZymoPURE II Plasmid Midiprep Kit (Zymo Research) or as linear expression templates (LETs). For LETs, eBlocks were inserted into pJL1 using Gibson Assembly with a linearized pJL1 backbone. Following Gibson Assembly (GA), each reaction was then diluted 10x in nuclease free water. 1 μL of diluted GA reaction was then used in a 50 μL PCR reaction using Q5 Hot Start High-Fidelity DNA Polymerase (New England Biolabs).

Linearized pJL1 backbone: gagcatcaaatgaaactgcaatttattcatatcaggattatcaataccatatttttgaaaaagccgtttctgtaatgaaggagaaaactcaccgaggcagttccataggatggcaagatcctggtatcggtctgcgattccgactcgtccaacatcaatacaacctattaatttcccctcgtcaaaaataaggttatcaagtgagaaatcaccatgagtgacgactgaatccggtgagaatggcaaaagcttatgcatttctttccagacttgttcaacaggccagccattacgctcgtcatcaaaatcactcgcatcaaccaaaccgttattcattcgtgattgcgcctgagcgagacgaaatacgcgatcgctgttaaaaggacaattacaaacaggaatcgaatgcaaccggcgcaggaacactgccagcgcatcaacaatattttcacctgaatcaggatattcttctaatacctggaatgctgttttcccggggatcgcagtggtgagtaaccatgcatcatcaggagtacggataaaatgcttgatggtcggaagaggcataaattccgtcagccagtttagtctgaccatctcatctgtaacatcattggcaacgctacctttgccatgtttcagaaacaactctggcgcatcgggcttcccatacaatcgatagattgtcgcacctgattgcccgacattatcgcgagcccatttatacccatataaatcagcatccatgttggaatttaatcgcggcttcgagcaagacgtttcccgttgaatatggctcataacaccccttgtattactgtttatgtaagcagacagttttattgttcatgatgatatatttttatcttgtgcaatgtaacatcagagattttgagacacaacgtgagatcaaaggatcttcttgagatcctttttttctgcgcgtaatctgctgcttgcaaacaaaaaaaccaccgctaccagcggtggtttgtttgccggatcaagagctaccaactctttttccgaaggtaactggcttcagcagagcgcagataccaaatactgttcttctagtgtagccgtagttaggccaccacttcaagaactctgtagcaccgcctacatacctcgctctgctaatcctgttaccagtggctgctgccagtggcgataagtcgtgtcttaccgggttggactcaagacgatagttaccggataaggcgcagcggtcgggctgaacggggggttcgtgcacacagcccagcttggagcgaacgacctacaccgaactgagatacctacagcgtgagctatgagaaagcgccacgcttcccgaagggagaaaggcggacaggtatccggtaagcggcagggtcggaacaggagagcgcacgagggagcttccagggggaaacgcctggtatctttatagtcctgtcgggtttcgccacctctgacttgagcgtcgatttttgtgatgctcgtcaggggggcggagcctatggaaaaacgccagcaacgcgatcccgcgaaattaatacgactcactatagggagaccacaacggtttc

Forward primer for LET PCR: ctgagatacctacagcgtgagc

Reverse primer for LET PCR: cgtcactcatggtgatttctcacttg

#### FluoroTect**^TM^** gel

PURE*frex* 2.1 (Gene Frontier) reactions were assembled according to manufacturer instructions, using 1 μL of unpurified template LET and 0.5 μL of FluoroTect^TM^ (Promega) per 10 μL reaction. Following incubation at 37 °C for 6 hours, samples were centrifuged at 12,000 xg for 10 min at 4 °C. 3 μL of supernatant was then mixed with 1 μL of 40 μg/mL RNase A and incubated at 37 °C for 10 minutes. Following incubation, 1 μL of 1M DTT, 2.5 μL of 4X Protein Sample Loading Buffer for Western Blots (Li-COR Biosciences), and 2.5 μL of water were added to each sample and the samples were then incubated at 70 °C for 10 minutes. Samples were then loaded on a NuPAGE 4-12% Bis-Tris Protein Gel and run for 40 minutes at 200 V in MES Running Buffer. For comparison, a lane was loaded with BenchMark fluorescent protein standard (Thermo Fisher Scientific). The resulting gel was then imaged using both the 600 and 700 fluorescent channel on a LICOR Odyssey Fc (Li-COR Biosciences).

#### AlphaLISA reactions for RiPPs

PURE*frex* 2.1 (Gene Frontier) reactions were assembled according to manufacturer instructions. Briefly, 1 μL of the unpurified LET reaction – encoding for the precursor peptide or RRE - was added as a template per 10 μL PURE*frex* reaction. Reactions were then incubated at 37 °C for 5 hours. After incubation, these samples were then diluted in a buffer consisting of 50 mM HEPES pH 7.4, 150 mM NaCl, 1 mg/mL BSA, and 0.015% v/v Triton X-100. Following dilution, an Echo 525 acoustic liquid handler was used to dispense 0.5 μL of diluted RRE, 0.5 μL of diluted peptide, and 0.5 μL of blank buffer from a 384-well polypropylene 2.0 Plus Source microplate (Labcyte) using the 384PP_Plus_GPSA fluid type into a ProxiPlate-384 Plus, White 384-shallow well destination microplate. The plate was then sealed and equilibrated at room temperature for one hour. Next, anti-FLAG Alpha Donor beads (Perkin Elmer) were used to immobilize the sFLAG tagged peptides and anti-Maltose-Binding (MBP) AlphaLISA acceptor beads were used to immobilize the MBP-tagged RREs. 0.5 μL of acceptor and donor beads diluted in buffer were added to each reaction to a final concentration of 0.08 mg/mL and 0.02 mg/mL donor and acceptor beads respectively. Reactions were then equilibrated an additional hour at room temperature in the dark. For analysis, reactions were incubated for 10 minutes in a Tecan Infinite M1000 Pro plate reader at room temperature and then chemiluminescence signal was read using the AlphaLISA filter with an excitation time of 100 ms, an integration time of 300 ms, and a settle time of 20 ms. Results were visualized using Prism version 9.5.1 (GraphPad).

#### Computational prediction of lasso peptide BGCs

A diverse collection of 39,311 publicly genomes available (2020 April) spanning soil bacteria, metagenomes and extremophiles were analyzed using AntiSMASH 5.1.2 identifying 315,876 biosynthetic gene clusters (**Supplementary Table 2**). A total of 2,574 lasso peptide clusters were identified, and from this set, 1,882 BGCs contained a complete collection of essential biosynthetic enzymes (**Supplementary Table 3**). Further prioritizing these clusters, a sequence similarity network^98,99^ of the identified propeptide genes with a collection of known lasso peptide sequences was created to assess the novelty of each cluster. Subsequent filtering of the remaining novel BGCs included selecting BGCs based on a propeptide length of 17-27 amino acids and whether the mature lasso peptide is predicted to carry a positive charge at a neutral pH. Calculation of the predicted isoelectric point of the predicted core peptides used Thermo Fisher Scientific’s peptide analysis tool (https://www.thermofisher.com/us/en/home/life-science/protein-biology/peptides-proteins/custom-peptide-synthesis-services/peptide-analyzing-tool.html). This narrowed the selection to 202 BGCs, of which 47 were chosen. A total of 210 genes were synthesized by Twist Bioscience. All amino acid sequences and metadata for the 47 selected BGCs are provided in the **Extended Data Set**.

#### *In vivo* expression and purification of lasso peptide tailoring enzymes

For computationally predicted MBP-RREs and MBP-proteases, constructs of the target protein in pET.BCS.RBSU.NS backbone were transformed into BL21 Star (DE3) cells, plated on LB agar plates containing 100 μg/mL carbenicillin, and incubated at 37 °C. Single colonies were cultured in 50 mL of LB containing 100 μg/mL carbenicillin at 37 °C and 250 RPM. After overnight incubation, 20 mL of the overnight culture were used to inoculate 1L of LB supplemented with 2 g/L glucose and 100 μg/mL carbenicillin. Cells were grown at 37 °C and 250 RPM and induced for protein production at OD_600_ 0.6-0.8 with 500 μL of 1M IPTG. Four hours post induction, cells were harvested via centrifugation at 5,000 xg for 10 minutes at 4 °C and flash frozen in liquid nitrogen.

After thawing on ice, cell pellets were resuspended in lysis buffer composed of 50 mM Tris-HCl pH 7.4, 500 mM NaCl, 2.5 % (v/v) glycerol, and 0.1% Triton X-100. For cell pellets used to overexpress RREs and cyclases, the lysis buffer also contained 6 mM PMSF, 100 μM Leupeptin, and 100 μM E64. Cell suspensions were then supplemented with 1 mg/mL lysozyme and lysed via sonication using a Qsonica sonicator at 50% amplitude for 2 minutes with 10 seconds on 10 second off cycles. Following sonication, insoluble debris were removed via centrifugation at 14,000 xg for 30 minutes at 4 °C. Per 1L of cell culture, 5 mL of amylose resin was equilibrated with 5 to 10 column volumes of wash buffer (50 mM Tris HCl, 500 mM NaCl, 2.5 % (v/v) glycerol, pH 7.4) in a 50 mL conical tube and mixed via inversion. Resin was separated from wash buffer by spinning at 2,000 xg for 2 min at 4 °C and the supernatant was then poured off. Equilibration was repeated for a total of 4 times with fresh equilibration buffer. Following the last equilibration, the cleared cell lysis supernatant was added to the resin and incubated for 2 hours at 4 °C with constant agitation on a shake table. Following incubation on the resin, the resin was washed once with 5 column volumes of lysis buffer followed by 5 column volumes of wash buffer four times. For the last wash, the resuspended resin was loaded in a 25 mL gravity flow column and drained via gravity flow. For elution, 15 mL of elution buffer (50 mM Tris HCl, 300 mM NaCl, 10 mM maltose, 2.5% (v/v) glycerol, pH 7.4) was added to the gravity flow column and collected. Samples were then buffer exchanged into storage buffer (50 mM HEPES, 300 mM NaCl, 0.5 mM TCEP, 2.5% (v/v) glycerol, pH 7.5) using amicon spin filters (50 kDa MWCO) by spinning at 4,500 xg for 10-15 minutes. Samples were then aliquoted, flash frozen, and stored at -80 °C until use. Total protein concentration of each purified sample was determined using a Bradford assay (Biorad). Percent purity of each sample was determined by running diluted aliquots of each purified protein on a 4-12% Bis-Tris gel and staining with Optiblot Blue (Abcam). After destaining, each gel was imaged using the 700 fluorescent channel on a LICOR Odyssey Fc (Li-COR Biosciences, USA) and percent purity was determined via densitometry using Licor Image Studio Lite. Final concentrations of each protein were then calculated by multiplying the total protein content by the percent purity.

Computationally identified cyclases were expressed and purified according to the process outlined above for computationally identified RREs except for transforming into BL21 Star (DE3) cells already transformed with pG-KJE8. LB agar and media for cell growth were supplemented with 20 μg/mL chloramphenicol in addition to 100 μg/mL carbenicillin. At inoculation, LB was supplemented with 2 g/L glucose, 100 μg/mL carbenicillin, 20 μg/mL chloramphenicol, and 2 ng/mL anhydrotetracycline per 1 L of media for induction of folding chaperones.

#### *In vitro* enzymatic assembly of lasso peptide BGCs

PURE*frex* 2.1 (Gene Frontier) reactions to express the precursor peptide were assembled according to manufacturer instructions using 1 μL of 200 ng/μL plasmid (pJL1 backbone encoding precursor peptide of interest) per 10 μL reaction and incubated at 37 °C for at least five hours. Purified proteins were buffer exchanged using Zeba Micro Spin Desalting Columns (7K MWCO) into synthetase buffer (50 mM Tris-HCl pH 7.5, 125 mM NaCl, 20 mM MgCl_2_). 10 μL reactions were then assembled using 5 μL of PURE*frex* reaction, and the appropriate volume of each individual purified enzyme or buffer such that both the RRE and protease were at a final concentration of 10 μM and the cyclase was at a final concentration of 1 μM. Reactions were supplemented to a final concentration of 10 mM DTT and 5 mM ATP and incubated at 37 °C for varying lengths of time. For analysis, samples were desalted using Pierce C18 spin tips (10 μL bed), spotted on a MALDI target plate using 50% saturated CHCA matrix in 80% ACN with 0.1% TFA, and analyzed using a Bruker RapiFlex MALDI-TOF mass spectrometer in reflector positive mode at Northwestern University’s Integrated Molecular Structure Education and Research Center (IMSERC).

#### Carboxypeptidase treatment of lasso peptides

Assembled reactions (20 μL scale) were desalted using Pierce C18 spin column and eluted into 20 μL of acetonitrile. After solvent removal under vacuum, reactions were resuspended in a solution containing carboxypeptidase Y at 50 ng/μL in 1X PBS (10 μL) and incubated at room temperature overnight. The mixtures were evaporated to dryness and resuspended in 3 μL saturated α-Cyano-4-hydroxycinnamic acid (CHCA) matrix solution in TFA (trifluoroacetic acid). Samples were then spotted on a matrix assisted laser desorption/ionization (MALDI) plate and analyzed using a Bruker RapiFlex MALDI-TOF mass spectrometer in reflector positive mode at Northwestern University’s Integrated Molecular Structure Education and Research Center (IMSERC).

### Methods for conjugate vaccine work

#### DNA design and preparation

For sequences used in the PglB mutant screen, the wild-type sequence for PglB was retrieved from Uniprot (Q5HTX9) and codon optimized using the IDT Codon Optimization Tool. A codon optimized linker and c-myc tag were appended to the C-terminus of the sequence. Each single variant sequence was then created using the codon optimized wild-type sequence as the template and replacing the respective codon with the most prevalent codon for the replacement amino acid. All peptide sequences were ordered as eBlocks with overhang to a linearized pJL1 backbone for use in Gibson Assembly reactions.

The wild-type sequence for *Haemophilus influenzae* protein D was retrieved from Uniprot (Q06282) and codon optimized using the IDT Codon Optimization Tool. Codon optimized linkers, a StrepII tag, and a 6xHis tag were appended to the C-terminus of the PD sequence. Each sequon variant was created by inserting the DNA sequence “AGAGCAGGAGGTGACCAGAACGCTACACGCGCAACCACA” (AA sequence: “RAGGDQNATRATT”) between each codon in the wild-type PD sequence. Eleven negative controls were added by instead inserting the DNA sequence “AGAGCAGGAGGTGACCAGTTGGCTACACGCGCAACCACA” (AA sequence: “RAGGDQLATRATT”), in which the asparagine in the sequon is replaced with a leucine that is not glycosylated. All sequon variants were ordered as eBlocks with overhang to linearized pJL1 backbone for use in Gibson Assembly reactions.

The cell-free library generation for the PglB mutant screen was prepared as follows: (1) each gBlock was amplified in a 50 µL PCR reaction with 0.1 ng template added; (2) PCR products were cleaned with a Clean and Concentrate kit (NEB); (3) cleaned PCR product was diluted to 6.7 ng/µL with nuclease-free water; (4) eBlocks were mixed with their respective pair of gBlocks for a final concentration of 1.5 ng/µL of each component in a 5 µL Gibson reaction Gibson reactions were performed to insert each PglB; (5) 4 µL of Gibson product was added to a 16 µL rolling circle amplification (RCA) reaction using phi 29-XT polymerase (NEB); (6) the completed RCA reaction was diluted 1:1 with the addition of 20 µL nuclease-free water. All PCR reactions used Q5 Hot Start DNA polymerase (NEB). The diluted RCA product serves as a template for expression of each PglB mutant in CFPS. To express sfGFP carrier protein, 200 µL CFPS reactions were prepared containing 13.3 ng/µL plasmid encoding the carrier.

The cell-free library generation for the PD sequon walking experiment was prepared using the same workflow as the PglB mutant screen, but with the following exceptions: (1) sequon variant eBlocks and backbone gBlocks were added to 5 µL Gibson reactions for a final concentration of 8 µM of each sequence; (2) Gibson reactions were diluted 6x in nuclease-free water; (3) 1 µL diluted Gibson product was added to 9 µL PCR reactions to generate linear expression templates of each sequon variant. Expression of each sequon variant was performed by adding 1 µL of linear expression template to a 4 µL CFPS reaction.

Forward primer for sequon variant LET PCR: CGATAGTTACCGGATAAGGC

Reverse primer for sequon variant LET PCR: CCATTCTCACCGGATTCAG

#### Cell Extract Preparation

Extract from BL21 StarTM (DE3) cells was prepared based on previous reports^100,101^. Briefly, an overnight culture was used to inoculate a culture of 2xYTPG at the 10 L scale (target optical density at 600 nm (OD600) = 0.06-0.08) in a Sartorius Stedim BIOSTAT Cplus bioreactor. The culture was then incubated at 37 °C with agitation set to 250 RPM. Once the culture reached OD600 = 0.6, the cells were induced for T7 RNA polymerase expression by adding IPTG to a final concentration of 0.5 mM. At OD600 = 3.0, the cells were harvested and centrifuged at 8,000 xg for 5 min. The resulting cell pellet was then collected and washed 3x with 25 mL of S30 buffer (10 mM Tris acetate pH 8.2, 14 mM magnesium acetate, and 60 mM potassium acetate) by vortexing in cycles of 15 seconds vortexing and 15 seconds on ice. In between each wash step, cells were pelleted via centrifugation at 10,000 xg for 2 min and the supernatant was poured off. After the final wash step, the supernatant was poured off, the mass of the cell pellet was recorded, and the cell pellets were flash frozen and stored at -80 °C. For lysate preparation, the cell pellets were thawed on ice for 1 hr. Next, 1 mL of S30 buffer per gram of cell pellet was added to each tube. The cells were then resuspended via vortexing, again in cycles of 15 seconds vortexing and 15 seconds on ice. After resuspension, the cells were then lysed via homogenization using a single pass through an Avestin EmulsiFlex-B15 homogenizer between 20,000-25,000 psig. Following homogenization, the lysed sample was centrifuged at 12,000 xg for 10 minutes at 4 °C. The supernatant was then collected and centrifuged again at 12,000 xg for 10 minutes at 4 °C. Following the final centrifugation, the supernatant was pooled, aliquoted, flash frozen, and stored at -80 °C until use.

For extracts enriched with *cj*PglB^Q287K^ and CPS from *S. pneumoniae* serotype 4 and derived from Hobby strain^82^, the above directions provided for BL21 Star^TM^ (DE3) cells were followed with the following changes: Prior to growing the overnight cultures, electrocompetent Hobby cells were transformed with pSF-*Cj*PglB^Q287K^-LpxE-KanR and pB-4^102^ and plated on LB agar plates containing 50 mg/mL Kanamycin and 20 mg/mL of tetracycline. During each cell growth phase, the cultures were also supplemented with 50 mg/mL Kanamycin and 20 μg/mL of tetracycline. At OD_600_ = 0.6-0.8, the culture was supplemented with 0.1% w/v arabinose in addition to 0.5 mM IPTG to induce for *cj*PglB^Q287K^ and CPS4 expression respectively, and the incubator was turned down to 220 RPM and 30 °C. Additionally, the supernatant from the first 12,000 xg centrifugation spin was collected and underwent runoff by wrapping the tubes in aluminum foil and incubating at 37 °C and 250 RPM for 1 hr. Following runoff, the tubes were centrifuged at 10,000 xg at 4 °C for 10 min and the supernatant was collected, mixed, and aliquoted before flash freezing and storing at -80 °C until use.

#### Crude membrane fraction

For producing crude membrane fraction, Hobby strain cells were transformed with pB-4^102^ and plated on LB agar plates containing 20 μg/mL of tetracycline. A single colony was then used to inoculate a 50 mL overnight culture of LB supplemented with 20 μg/mL of tetracycline. The next morning, 1 L of 2xYTPG supplemented with 20 μg/mL of tetracycline was inoculated with a target starting OD_600_ = 0.06-0.08. The culture was the incubated at 37 °C with agitation set to 250 RPM. At OD_600_ = 0.6-0.8, the culture was supplemented with 0.5 mM IPTG and the culture was then incubated overnight at 30 °C and agitation of 220 RPM. The next morning, the cells were harvested via centrifugation at 8,000 xg for 5 min at 4 °C. After pouring off the supernatant, 1 mL per gram of cell pellet of resuspension buffer (50 mM Tris HCl, pH 7.5, 25 mM NaCl) was added to the pellets. The cells were then resuspended via vortexing in cycles of 15 second vortexing and 15 second on ice. After the cells were fully resuspended, the sample was lysed via homogenization using a single pass through an Avestin EmulsiFlex-B15 homogenizer between 20,000-25,000 psig. Following lysis, the sample was then centrifuged at 12,000 xg at 4 °C for 30 minutes. The supernatant was then ultracentrifuged at 100,000 xg at 4 °C for 1 hour to pellet the membrane vesicles. Following ultracentrifugation, the supernatant was poured off and 0.2 mL/gram original cell pellet of resuspension buffer (50 mM Tris HCl, pH 7.5, 25 mM NaCl, 1% w/v DDM) was added to the pellet before incubating overnight at 4 °C on a shake table. The next morning, the samples were pipette mixed to ensure complete resuspension of the pellet and the incubated at room temperature for 30 min. Finally, the samples were spun at 16,000 xg for 1 hour at 4 °C and the supernatant was mixed, aliquoted, flash frozen, and stored at -80 °C until use.

#### Nanodisc supplemented CFPS reactions

*In vitro* expression of each WT or mutant PglB construct was performed by adding 1 µL of diluted RCA product to a 4 µL BL21 Star^TM^ (DE3) CFPS reaction supplemented with 66.7 µM MSP1E3D1 POPC nanodiscs (Cube Biotech).

#### *In vitro* glycosylation reactions

In the PglB mutagenesis screen, *in vitro* glycosylation reactions were performed by combining 0.4 µL unpurified sfGFP acceptor, 1 µL unpurified PglB mutant, and 3 µL S. pneumoniae CPS 4 crude membrane fraction in a 5 µL reaction volume containing 0.1% w/v DDM (Anatrace), 1% w/v Ficoll 400 (Sigma), 10 mM manganese chloride (Sigma), and 50 mM HEPES (Sigma).

In the sequon walking experiment, *in vitro* glycosylation reactions were performed by combining 1 µL unpurified sequon variant and 2.5 µL enriched extract containing PglB^Q287K^ and S. pneumoniae CPS 4 in a 5 µL reaction volume containing 0.1% w/v DDM, 1% w/v Ficoll 400, 10 mM manganese chloride, and 50 mM HEPES.

#### AlphaLISA reactions for glycoconjugates

Completed *in vitro* glycosylation reactions were diluted in a buffer consisting of 50 mM HEPES pH 7.4, 150 mM NaCl, 1 mg/mL BSA, and 0.015% v/v Triton X-100. All glycoconjugate AlphaLISA experiments were performed with 1 µL reaction volumes with a 0.08 mg/mL final concentration of Protein A donor beads and 0.02 mg/mL final concentration of anti-6xHis acceptor beads, which immobilize the *S. pneumoniae* CPS 4 anti-serum and the 6xHis-tagged glycoconjugates, respectively. Following dilution, an Echo 525 acoustic liquid handler was used to dispense 0.25 µL diluted *in vitro* glycosylation product, 0.25 µL *S. pneumoniae* CPS 4 anti-serum, 0.25 µL blank buffer, and 0.125 µL anti-6xHis acceptor beads diluted in buffer from a 384-well polypropylene 2.0 Plus Source microplate using the 384PP_Plus_GPSA fluid type into an AlphaPlate 1536-well destination microplate (Revvity). The plate was sealed and equilibrated for one hour at room temperature. Following incubation, 0.125 µL of Protein A donor beads diluted in buffer were transferred to each reaction. Reactions were equilibrated for an additional hour at room temperature in the dark. For analysis, reactions were incubated for 10 minutes in a Biotek Synergy Neo2 plate reader at room temperature, and chemiluminescent signal was read using the AlphaLISA filter with an excitation time of 100 ms, an integration time of 300 ms, and a settle time of 20 ms. For the PD sequon walking experiment, replicate AlphaLISA reactions were performed on separate plates. Signal for each plate was normalized using the formula: Normalized signal = (Raw signal – mean neg. control signal)/mean pos. control signal. Results were visualized using Prism version 10.3.1 (GraphPad).

#### Western blotting

Samples were loaded on a 4-12% Bis-Tris gel and run with either MOPS SDS or MES SDS buffer for 45 min at 200V. A semidry transfer cell was then used to transfer the samples to Immobilon-P-poly(vinylidene difluoride) PVDF 0.45 μm membranes at 80 mA per blot for 45 min. After transferring, the membranes were blocked for 30 min at room temperature in intercept blocking buffer with gentle shaking. Following blocking, the blots were briefly rinsed with 1x PBST and then probed for 1 hour at room temperature with gentle shaking using one of the following antibodies diluted into intercept binding buffer with 0.2% Tween20: anti-6xHis (Abcam, ab1187) at 1:7500 dilution, anti-type 4 pneumococcal antiserum (Cedarlane, 16747(SS)) at 1:1000 dilution, or anti-myc (Abcam, ab9106) at 1:000 dilution. Following primary incubation, membranes were rinsed twice with 1x PBST followed by 3 five min washes in 1x PBST at room temperature with gentle shaking. Following washing, the blots were probed for 1 hour at room temperature with gentle shaking using a fluorescent goat, anti-rabbit antibody GAR-680RD (Licor, 926-68071) at a dilution of 1:10,000 in intercept blocking buffer with 0.2% Tween20 and 0.1% SDS. Then, the membranes were washed as described earlier after probing with the primary antibody. Finally, the blots were imaged with either Licor Image Studio or an Azure 600 imager and analyzed by densitometry using Licor Image Studio Lite. The fluorescence background was subtracted from each membrane before assessing densitometry.

## Acknowledgements

The authors would like to thank Rui Gan, Jonathan Bogart, and Thuy Aziz for helpful discussions. This work was supported by the National Institutes of Health (NIH) 1U19AI142780-01. D.A.W. acknowledges support from the National Science Foundation Graduate Research Fellowship under grant no. DGE-1842165. Z.M.S. acknowledges support from the National Science Foundation National Research Traineeship under grant no. 2021900. M.D. acknowledges support from the Canadian Institutes of Health Research Postdoctoral Fellowship under grant no. MFE-176575. This work made use of the IMSERC MS facility at Northwestern University, which has received support from the Soft and Hybrid Nanotechnology Experimental (SHyNE) Resource (NSF ECCS-2025633), the State of Illinois, and the International Institute for Nanotechnology (IIN).

## Author Contributions

D.A.W. designed research, performed experiments, performed MALDI-MS on reactions, analyzed data, and wrote the paper. Z.M.S. designed research, performed experiments, analyzed data, and wrote the paper. M.D.C. designed research, performed experiments, performed MALDI-MS on reactions, analyzed data, and edited the paper. M.D. computationally identified all lasso peptide BGCs and wrote the paper. K.F.W performed experiments. D.V.P. performed experiments. S.E.S. performed experiments. R.F. performed experiments. R.N. supervised research and edited the paper. M.P.D supervised research and edited the paper. E.P.B. supervised research and edited the paper. A.S.K. supervised research, analyzed data, and edited the paper. M.C.J. designed and directed research, analyzed data, and wrote the paper.

